# Dominance of plasticity in wind-pollinated trees’ flowering time response to temperature

**DOI:** 10.1101/2025.08.26.672405

**Authors:** Yi Liu, Yiluan Song, William Weaver, Kerby Shedden, Yang Chen, Stephen Smith, Kai Zhu

## Abstract

Climate change is shifting the flowering phenology of wind-pollinated trees, leading to longer pollen seasons and heightened allergy risks. Accurately forecasting these phenological shifts requires a clear understanding of the underlying mechanisms and variations in the sensitivity of phenology to temperature. Here, we investigated the temperature sensitivity of flowering phenology in 74 wind-pollinated tree species across the US using long-term herbarium records spanning over a century. We found consistent phenological responses to spring temperature variations over space and over time in 77% of species, indicating that these long-lived trees can rely primarily on plasticity to cope with temperature variation across different regions. Sensitivity estimates were consistent across both century-scale and decadal time scales, challenging the previous understanding of declines. Interspecific variations in sensitivity exhibited a strong phylogenetic signal, indicating evolutionary conservatism in phenological responses among related species. Together, these findings shed light on the magnitude, mechanism, trend, and variability of temperature sensitivity of wind-pollinated tree flowering phenology. Our results improve the mechanistic basis for forecasting phenological shifts and their downstream effects on airborne pollen and allergen exposure under ongoing climate change.

## 1. Introduction

Climate-driven shifts in the flowering phenology of wind-pollinated plants are lengthening pollen seasons and increasing allergy risks. Recent studies show that climate change has contributed to a 20-day extension of the pollen season and a 21% increase in pollen concentrations since 1990 (Anderegg et al., 2021), which might exacerbate allergic respiratory conditions, such as hay fever and asthma (Ng & Boersma, 2023; Zablotsky et al., 2023). At the root of these changes in pollen dynamics lies the responses of flowering phenology in wind-pollinated plants, which are the primary source of airborne pollen (Lo et al., 2019). Wind-pollinated plants’ flowering phenology has been reported to advance at a rate of 3−5 days per decade in response to recent climate trends (Ziello et al., 2012). An important quantity for evaluating and predicting these changes is *sensitivity*, defined as the change in timing of phenological events per unit change in an environmental factor (Wolkovich et al., 2021). Temperature sensitivity of flowering phenology in wind-pollinated species provides critical information for predicting shifts in flowering time under projected warming and anticipating resulting changes in pollen seasons.

Despite widespread recognition of the importance of temperature sensitivity, reported sensitivities vary considerably across studies. Meta-analyses of the literature and analyses of herbarium data indicate that temperature sensitivity can vary widely, from advancing by 10 days per °C to delaying by 5 days per °C (Fitter & Fitter, 2002; Menzel et al., 2006; Ramirez-Parada et al., 2024). Different estimates may lead to significant under- and overestimation of plant phenological responses to climate change (Wolkovich et al., 2012). These inconsistencies might reflect differences in the mechanistic focus of study (plasticity or adaptation) (Knott et al., 2023; Valdés et al., 2023), the temporal scale of responses (interannual or decadal) (Wolkovich et al., 2012), as well as phenological strategies of the taxa (Xie et al., 2022).

Disentangling these three sources of variation in temperature sensitivity is critical for predicting how different wind-pollinated species will respond to climate change, in both the short and long terms, through various biological processes.

First, variations in flowering phenology arise from two primary mechanisms: phenotypic plasticity and local adaptation, shown in observational studies and common garden experiments (Knott et al., 2023; Valdés et al., 2023). Phenotypic plasticity is the ability of an individual organism to adjust its physiology, development, or behavior in response to environmental changes without genetic alteration. In contrast, local adaptation refers to long-term genetic differentiation that accumulates across generations within a population, leading to traits that increase fitness in a specific environment (Fox et al., 2019). Short-lived species can evolve rapidly through genetic changes that enhance adaptation to their environment, whereas long-lived species evolve more slowly and may depend more heavily on broad phenotypic plasticity to cope with temperature variation (Cordero & Janzen, 2013). These two mechanisms are often reflected implicitly in the estimation of temperature sensitivity. *Temporal sensitivity* quantifies interannual variation at a single site, reflecting phenotypic plasticity. *Spatial sensitivity*, by contrast, quantifies variation across sites. If driven by plasticity alone, spatial sensitivity should equal temporal sensitivity because temperature differences of equivalent magnitude should elicit identical plastic responses in flowering time, whether expressed spatially (across sites) or temporally (across years). Otherwise, a difference between spatial and temporal sensitivity indicates local adaptation, with the direction and magnitude of this difference revealing the nature and extent of adaptation (Ramirez-Parada et al. 2024). Despite the distinct mechanisms, a common practice in global change research is to substitute spatial sensitivity for temporal sensitivity when predicting future changes, due to the scarcity of long-term records, a method known as “space-for-time substitution” (Blois et al., 2013; Knapp et al., 2024; Knott et al., 2023; Perret et al., 2024). While practical, this approach can be misleading if applied without validation (Evans et al., 2025); for example, studies using urban heat islands as warming proxies often underpredict the temporal sensitivity of phenology (Blumstein et al., 2025). To ensure reliable projections, it is essential to test the validity of space-for-time substitutions by assessing the role of plasticity on shaping the spatial and temporal flowering time variation.

Second, temperature sensitivity might change over time due to a combination of environmental constraints, nonlinear response, and adaptation, making long-term predictions based on short-term observations or experiments unreliable. Studies have shown that the temperature sensitivity of spring phenology has declined in recent years, possibly due to limitations from other environmental cues such as chilling, photoperiod (Fu et al., 2015), or precipitation (Xiong et al., 2024). With continued warming, the influence of each additional degree of temperature rise on flowering advancement may weaken, leading to a decline in observed linear sensitivity over time (Wolkovich et al., 2021). Moreover, the underlying drivers of temperature sensitivity might shift, transitioning from being primarily governed by plastic responses to involving both plasticity and local adaptation, thereby altering phenological responses to climate change (Wolkovich et al., 2012). These changes in temperature sensitivity over time can cause long-term phenological responses to diverge from predictions based on short-term observations (Fu et al., 2015; Xiong et al., 2024) or experimental results (Wolkovich et al., 2012). Understanding these dynamics is critical for evaluating the reliability of long-term phenological forecasts and determining whether predictions should instead be limited to shorter time horizons due to changing sensitivities.

Finally, flowering responses to temperature vary across plant species, yet the extent and patterns of these variations are not well understood for wind-pollinated species. For insect-pollinated plant species, species-specific responses in flowering phenology have been suggested to disrupt the ecological synchrony in flowering among species and negatively impact pollination (Kudo & Ida, 2013; Xie et al., 2022). A similar understanding is needed for wind-pollinated plant species in order to anticipate possible “shuffling” of flowering time across species (Roberts et al., 2015), cascading to the pollen season. Beyond idiosyncratic variations among species, lineages may differ systematically in their capacity to track climate change through phenological shifts. Closely related species tend to exhibit similar flowering and leaf-out responses to temperature cues, likely reflecting shared genetic constraints and conserved developmental pathways (Davies et al., 2013). Incorporating phylogenetic information has been shown to improve species-level phenology forecasts (Morales-Castilla et al., 2024). However, the strength of the phylogenetic signal in flowering time shifts of wind-pollinated trees remains unknown. Understanding this signal is particularly important for predicting climate-driven flowering responses in wind-pollinated species, especially in this case, where many of them are understudied or undersampled.

In this study, we aim to address these three knowledge gaps in our understanding of temperature sensitivity in the flowering phenology of wind-pollinated trees by investigating the contributions of plasticity and adaptation, the differences across temporal scales, and the magnitude and structure of interspecific variations. We leverage two large and complementary data sources: herbarium records provide phenological observations spanning over a century, and a citizen science program that has amassed temporally dense field observation data over the past two decades. Using a combination of these spatiotemporal data sources, along with phylogenetically informed models, we ask three questions regarding the main sources of variation in temperature sensitivity of wind-pollinated trees: (1) To what extent are the sensitivities driven by phenotypic plasticity alone? (2) How do the sensitivities differ when quantified on different time scales? (3) How much do the sensitivities vary by species, and how are these variations explained by phylogenetics? By understanding the roles of plasticity, adaptation, temporal scale, and phylogeny in shaping the temperature sensitivity of wind-pollinated trees’ flowering phenology, our study represents a critical step in improving the ecological forecasting of shifting pollen seasons under climate change.

## 2. Methods

### 2.1. Assessing the role of plasticity by contrasting spatial and temporal sensitivities

To quantify the contributions of phenotypic plasticity to the responses of flowering phenology to spring temperature, we adapted the framework of Ramirez-Parada et al. (2024) that compares the magnitudes of spatial and temporal sensitivity. Temporal sensitivity quantifies phenological variations across years, driven by fluctuations in temperature, reflecting the role of plasticity. Spatial sensitivity quantifies phenological variations across sites due to differences in long-term average temperature, reflecting the combined effects of plasticity and adaptation. If phenotypic plasticity is the sole driver of variation, a given change in temperature is expected to produce a consistent shift in flowering time, regardless of whether the temperature variation occurs temporally (across years) or spatially (across locations).

Consequently, spatial and temporal sensitivities should be equivalent. In contrast, a divergence between spatial and temporal sensitivity suggests the influence and magnitude of local adaptation. To investigate this, we extracted flowering time data from herbarium specimens and integrated these records with corresponding climate data to quantify both spatial and temporal sensitivities and statistically test their differences.

#### 2.1.1 Flowering time data from herbarium specimens

In order to estimate sensitivity over space and time, we derived a novel dataset of flowering time observations using digitized herbarium specimen images from the Global Biodiversity Information Facility (GBIF). GBIF, which traditionally focuses on species occurrence data, also provides valuable phenological information on a broad spatial and temporal scale, particularly through its digitized herbarium specimens (Ramirez-Parada et al., 2024). Herbarium specimens are preserved plants that are dried and mounted on a sheet of paper with labels documenting collection details such as the collector’s name, collection date, and location (Daru, 2025). For wind-pollinated trees, herbarium specimens typically consist of branches, and the presence or absence of flowers on the branch indicates whether or not the collection time is within the short flowering period (Supplementary Figure 1). We focused on the seven most allergenic wind-pollinated angiosperm genera (Lo et al., 2019) and downloaded 259,293 images and metadata of specimens that were georeferenced, and obtained precise collection dates, collected within the conterminous United States from 1895 to 2023 (GBIF, 2024).

We used LeafMachine2 (Weaver & Smith, 2023), a computer vision tool, to identify specimen images with flowers. LeafMachine2 employs a trained object detection network to detect and classify image components into categories such as leaves, fruits, flowers, and buds while assigning confidence scores (Supplementary Figure 1). We summarized the results for each image based on the number of detected segments in each category with a confidence score greater than 0.2. A specimen was categorized as “flowering” if both a single flower and a flower cluster were detected, thereby minimizing false positives. The collection location was treated as the observational site, and the collection date was used as the flowering date. To preserve seasonal continuity between winter and spring, the flowering date was converted to “days of the year since November 1st.”

#### 2.1.2 Climate data

For all sites and years with valid flowering time, we retrieved monthly average temperature data at a 4 km spatial resolution from the Parameter–elevation Regressions on Independent Slopes Model (PRISM, http://prism.oregonstate.edu). PRISM was chosen as the source of climate data because it provides the longest available climate dataset, extending back to 1895. We calculated spring temperature as the mean monthly average temperature from March 1st to May 31st.

#### 2.1.3 Inferring the role of plasticity by comparing spatial and temporal sensitivity

We estimated spatial and temporal sensitivity by regressing flowering dates on the multi-year (1895−2023) average temperature (for spatial sensitivity) and the annual temperature anomaly relative to that average (for temporal sensitivity). We fitted a robust linear regression (Supplementary Method 1) model for each species using the *MASS* package in R (Figure 1b & c):

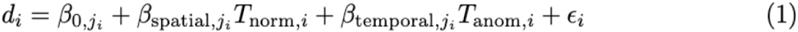

**Figure 1.**
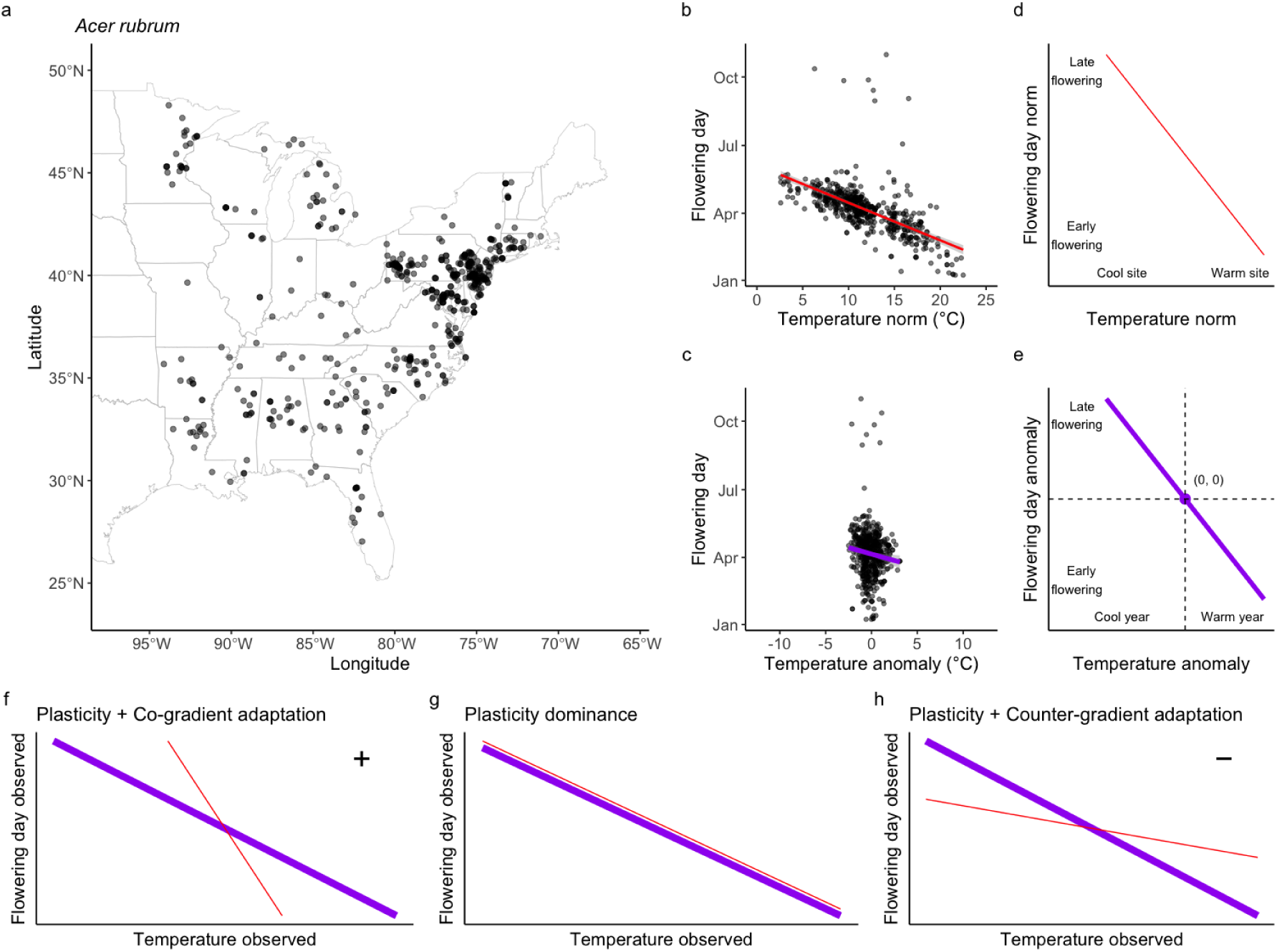
Assessing the role of plasticity in flowering’s response to temperature through calculating spatial and temporal sensitivity. **(a)** The spatial distribution of flowering time observations from herbarium data, using the most common tree species in the US, red maple (*Acer rubrum*), as an example. **(b)** Flowering time’s change with temperature norm, inferring spatial sensitivity. **(c)** Flowering time’s change with temperature anomaly, inferring temporal sensitivity. **(d)** Conceptual figure for spatial sensitivity: how many days of flowering time will advance in a location with a 1 °C increase in temperature. **(e)** Conceptual figure for temporal sensitivity: how many days of flowering time will advance in a year with a 1 °C increase in temperature. **(f)** The case of co-gradient adaptation (“+” symbol), where temporal sensitivity is greater than spatial sensitivity, indicating adaptation is in the same direction as and enhances plasticity. **(g)** The case where no significant differences between temporal sensitivity and spatial sensitivity, indicating the dominance of plasticity. **(h)** The case of counter-gradient adaptation (“−” symbol), where temporal sensitivity is weaker than spatial sensitivity, indicating adaptation is in the opposite direction and reduces plasticity.

where *i* is an observation indicator and *j_i_* is the species indicator, giving the species *j* to which observation *i* belongs. *d_i_* represents the day of flowering. *T*_norm,*i*_ is the multi-year average temperature (“norm”), which is determined by the location (not year) of the observation. *T*_anom,*i*_ is the temperature anomaly (deviation from *T*_norm,*i*_), which is determined by both the location and the year of the observation. The regression coefficient, spatial sensitivity *β*_spatial_, measures how flowering time varies across locations with differing long-term average temperatures *T*_norm_ (Figure 1b & d). The coefficient, temporal sensitivity *β*_temporal_, quantifies how flowering time responds to interannual temperature anomalies *T*_anom_ (Figure 1c & e). We retained only species with observations spanning at least 10 different *T*_norm_ values and 10 different *T*_norm_ values, with a minimum of 30 total observations, resulting in 10,482 flowering specimens from 74 species (Supplementary Figure 2).

The difference between spatial and temporal sensitivity reveals whether plasticity can explain the flowering variation alone (Ramirez-Parada et al., 2024). Dominated by plasticity, equivalent temperature differences are expected to drive the same plastic responses in flowering time, regardless of whether those differences occur across sites or across years. As a result, spatial and temporal sensitivities should be equal when only phenotypic plasticity is involved (Figure 1g). Co-gradient adaptation (Figure 1f) occurs when spatial sensitivity is stronger (more negative) than temporal sensitivity. This suggests that local adaptation amplifies the plastic response, with populations in warmer sites flowering even earlier than expected based on plasticity alone. Counter-gradient adaptation (Figure 1h) is observed when spatial sensitivity is weaker (less negative) than temporal sensitivity. Here, local adaptation counteracts the plastic response, resulting in populations in warmer sites flowering later than plasticity would predict. We used the Wald test (see Supplementary Method 2) to statistically compare the estimated *β*_temporal_ and *β*_spatial_ with their uncertainties. If the test indicates no significant difference, we conclude no evidence of local adaptation (Figure 1g). A significantly stronger spatial sensitivity indicates co-gradient adaptation, while a significantly weaker spatial sensitivity indicates counter-gradient adaptation.

### 2.2. Comparing long- and short-term sensitivities

Herbarium data offer flowering phenology information with broad temporal coverage spanning the past century. However, these records are temporally sparse, making it difficult to infer temperature sensitivity in recent years. To assess whether the temperature sensitivity of flowering phenology has changed in recent years, we used recent temporally dense field observation data to estimate sensitivities and compared these results to sensitivity estimates derived from long-term, temporally sparse herbarium records.

#### 2.2.1 Flowering time data from field observation

To estimate short-term temperature sensitivity with more data, we supplemented field observation data from the National Phenology Network (NPN), which has more data in the past two decades. NPN is a national-scale field monitoring system for phenology. It compiles data across the US since 2007 contributed by both professional scientists (e.g., NEON phenology plots) and community scientists (e.g., individual volunteers and organizations) using standardized protocols through platforms such as Nature’s Notebook (Rosemartin et al., 2018). Observers contributed raw data in the form of “yes” or “no” observations, indicating whether a plant was identified to be in a given phenophase (e.g., flowering) on a specific day. These records were processed by NPN into individual-level phenometrics, indicating the day of year with the first “yes” observation for a phenophase for each individual plant in each year. We used the individual-level phenometrics data for the “open flowers” phenophase as the flowering date. To constrain the uncertainty of the flowering date, we excluded observations with conflict between observers and required at least one “no” observation within 7 days before the first “yes” observation. We also transformed phenology into days since November 1st, as in the herbarium dataset.

#### 2.2.2 Inferring short- and long-term sensitivity change

We fitted the same species-specific models (Equation 1) to observations from the NPN field observation dataset, which are on a shorter temporal scale compared to the herbarium dataset. Then we examined whether there was an overlap in the 95% confidence interval for sensitivities from the two datasets by species. Since the sensitivity estimates from the two datasets were derived independently, the lack of overlap in their confidence intervals indicates, with 99% confidence level, that the estimates are significantly different. This suggests a significant difference in responses between the last century and the most recent two decades. If the sensitivity from short-term field observations is significantly lower (i.e., a stronger advancement in phenology in response to warming) than that from long-term herbarium data, it implies that sensitivity has increased in the recent two decades relative to the past century (denoted by a “+” symbol). Conversely, if the sensitivity from short-term observations is significantly higher (i.e., a weaker advancement or a delay), it indicates a decline in sensitivity in recent decades (denoted by a “–” symbol).

### 2.3. Inferring the pattern of sensitivity variation among species

Sensitivities vary among species, but this variation may not be entirely random or independent. It could follow a normal distribution or reflect phylogenetic relationships. To infer the interspecific variation and explore the potential influence of phylogeny, we fitted two hierarchical models, including a standard mixed-effects model (SMM) and a phylogenetic mixed-effects model (PMM).

#### 2.3.2 Modelling the across-species sensitivity variation

We first constructed a standard mixed-effects model to represent interspecific variation in sensitivity as a normal distribution, allowing us to infer the degree of convergence in sensitivity across species.

Compared to the species-specific model (Equation 1), where the parameters of each species are completely independent, the standard mixed-effects model (Equation 2) allows information sharing across all species, assuming that the species have similarity in their phenological response to temperature to some extent. Specifically, the parameters for each species were considered to be independently drawn from normal distributions.

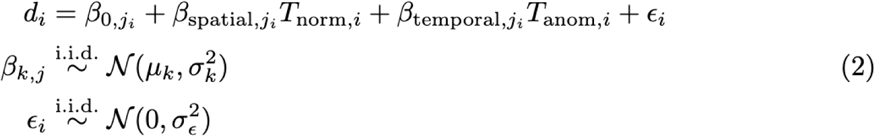

Here, *β_k_* represents a set of parameters including *β*_0_, *β*_spatial_, and *β*_temporal_. The hyperparameters *μ_k_* and *σ_k_*define the distributions from which each species-level parameter *β_k_*is drawn. We used RStan to fit the standard mixed-effects model; details of the model fitting are provided in the Supplementary Method 3.

#### 2.3.1 Phylogenetic data

Evolutionary history may shape temperature sensitivity variation, so we retrieved phylogenetic data representing evolutionary relationships among species. Phylogenetic data is typically in tree format, consisting of branches representing evolutionary lineages and nodes representing speciation events or shared ancestry. We employed the phylogenetic data for seed plants published by Smith & Brown (2018) using the phylo.maker function in the *V.PhyloMaker2* package (Jin & Qian, 2022). The phylo.maker function creates phylogenetic trees in three different ways. We used scenario S1, where any genera or species missing from the main tree are added at the base of their family. This is the most conservative approach because it avoids guessing where missing taxa belong within their families.

#### 2.3.3 Modelling the across-species sensitivity variation with phylogeny

To explore how lineages may differ systematically for temperature sensitivity, we further fitted a phylogenetic mixed-effects model (phylogenetic mixed-effects model), which incorporated phylogenetic data (Equation 3). Compared to the standard mixed-effects model that assumes normal distributions of parameters among species (Equation 2), the phylogenetic mixed-effects model uses multivariate normal distributions that account for phylogenetic distance between species (*ρ*) and the strength of phylogenetic signal (Pagel’s *λ*) when inferring how species share information. This approach assumes that species are not independent; those with closer phylogenetic relationships exhibit more correlated sensitivities.

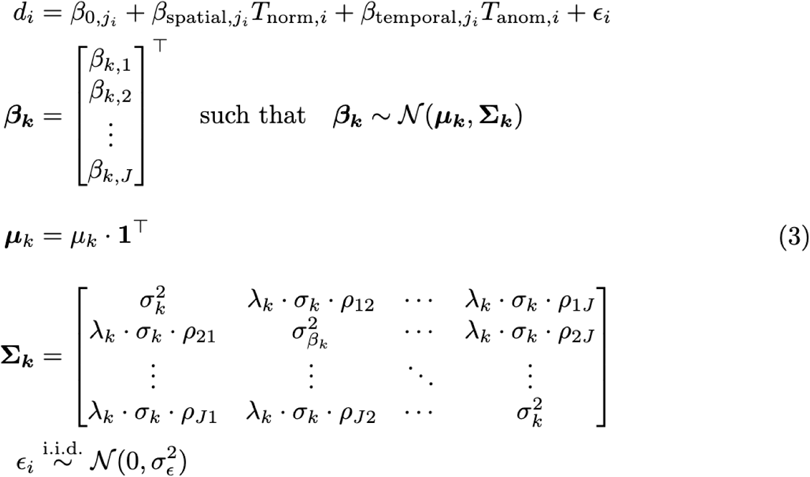

Here, each species-specific coefficient *β_k,j_* can be written in a vector ***β_k_***, which was modeled from a multivariate normal distribution with a mean vector ***μ****_k_* (all elements equal to *μ_k_*) and a structured covariance matrix **Σ***_k_*. This covariance matrix incorporated three key components: standard deviation (*σ_k_*), phylogenetic distance (*ρ*), and Pagel’s *λ* (*λ_k_*). *σ_k_*is the shared marginal standard deviation. *ρ* is the phylogenetic distance defined as the length of shared evolutionary history between species pairs (from their common ancestor to speciation). As an input parameter in the model, *ρ* determines pairwise correlations such that phylogenetically closer species exhibit stronger parameter correlations. *λ_k_* is a scaling factor representing phylogenetic signal strength (model output). Pagel’s *λ* is between 0 and 1, representing whether traits are shaped by evolutionary constraints (closer to 1) or random processes (closer to 0). In other words, a phylogenetic mixed-effects model with *λ* = 0 is equivalent to a standard mixed-effects model. We again used RStan to fit a phylogenetic mixed-effects model, and full details of model-fitting procedures are provided in the Supplementary Method 3.

## 3. Results

### 3.1 Dominance of plasticity given consistent spatial and temporal sensitivities

For most species, we found consistent temporal and spatial sensitivity based on herbarium specimen-derived flowering phenology, indicating that temperature sensitivity is dominated by phenotypic plasticity (Figure 2, Table 1). Spatial sensitivities have a mean value of −4.89 days °C^−1^ with a standard deviation of 3.06 days °C^−1^. Temporal sensitivities have a mean value of −3.68 days °C^−1^ with a standard deviation of 3.96 days °C^−1^. Across the 74 species from the seven genera analyzed, 57 species (77%) showed no significant differences between spatial and temporal sensitivity, indicating the dominance of plasticity. *Morus* (100%, 3 species) and *Betula* (90%, 9 species) have the highest percentage of species showing dominance of plasticity. Nevertheless, there was a tendency of co-gradient adaptation, with 50 species (68%) showing spatial sensitivity stronger than temporal sensitivity, but only 12 of them were significant. *Fraxinus* showed the highest percentage (43%, 3 species) of species with significant adaptation, with one species (*F. dipetala*) showing co-gradient adaptation and two species (*F. cuspidata* and *F. latifolia*) showing counter-gradient adaptation.

**Figure 2.**
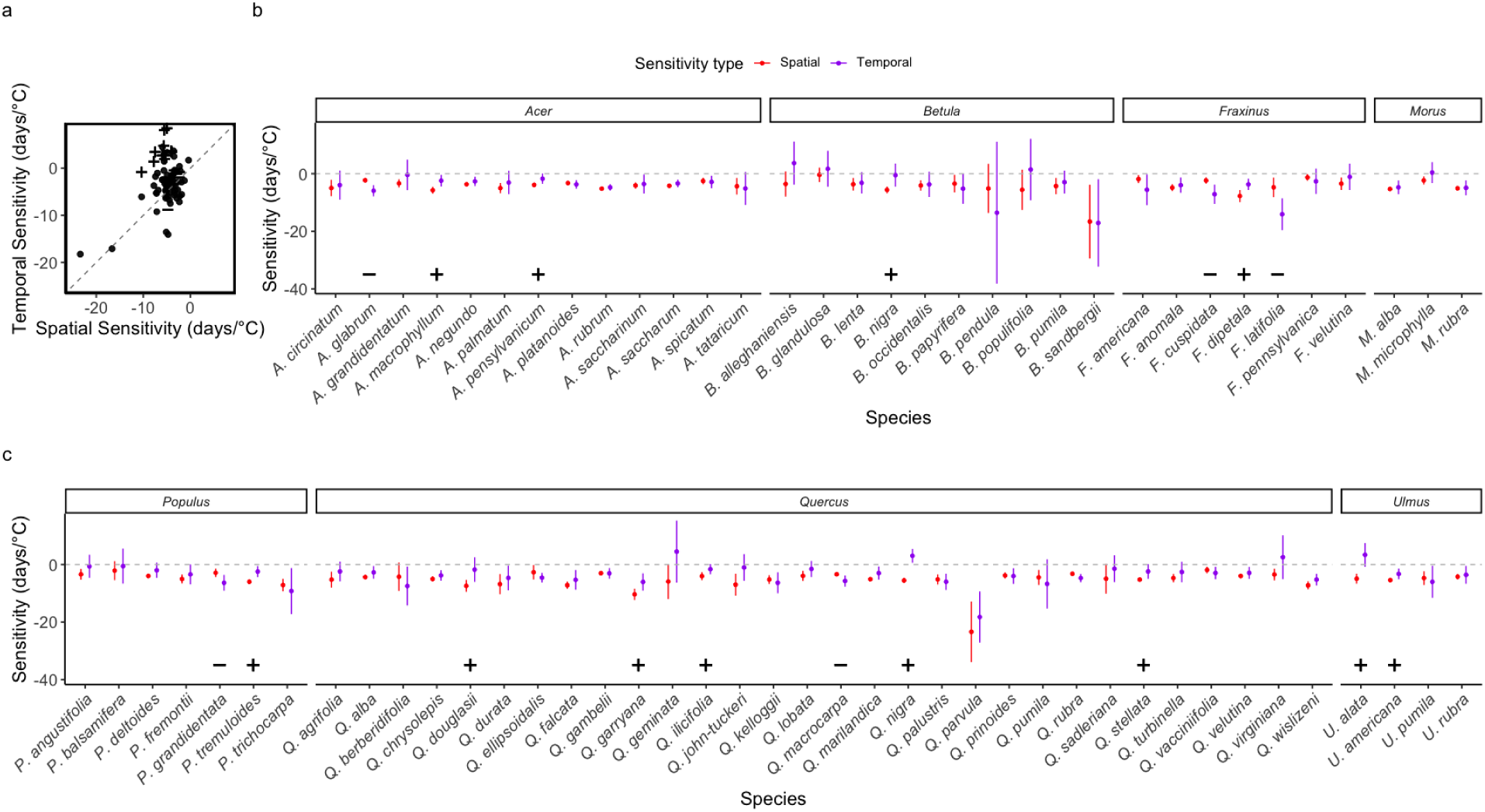
Comparison of spatial and temporal sensitivity from the species-independent model. **(a)** Scatter plot of spatial and temporal sensitivity of each species, with a grey dashed line representing the 1:1 line. Symbol “+” represents a significant co-gradient adaptation, and “−” represents a significant counter-gradient adaptation. Dots represent no significant adaptation. **(b) & (c)** By species spatial and temporal sensitivity comparison with the dot representing the point estimate and the line representing the 95% confidence interval. Symbol “+” represents a significant co-gradient adaptation, and “−” represents a significant counter-gradient adaptation.

**Table 1.** By genus summary of the plasticity role.

| Genus | Species number | Plasticity dominance | Counter-gradient adaptation | Co-gradient adaptation |
| --- | --- | --- | --- | --- |
| *Acer* | 13 | 10 (77%) | 1 (8%) | 2 (15%) |
| *Betula* | 10 | 9 (90%) | 0 (0%) | 1 (10%) |
| *Fraxinus* | 7 | 4 (57%) | 2 (3%) | 1 (14%) |
| *Morus* | 3 | 3 (100%) | 0 (0%) | 0 (0%) |
| *Populus* | 7 | 5 (71%) | 1 (1%) | 1 (14%) |
| *Quercus* | 30 | 24 (80%) | 1 (1%) | 5 (17%) |
| *Ulmus* | 4 | 2 (50%) | 0 (0%) | 2 (50%) |
| Total | 74 | 57 (77%) | 5 (7%) | 12 (16%) |

### 3.2 Consistent spatial and temporal sensitivities at different time scales

The long-term spatial and temporal sensitivities inferred from herbarium data collected over 1895-2022 aligned with the short-term sensitivities inferred from field observations collected over 2007-2023 (Figure 3, Table S1 & 2). Herbarium estimated sensitivities of −4.18 days °C^−1^ with a standard deviation of 1.18 days °C^−1^ (spatial) and −3.12 days °C^−1^ with a standard deviation of 1.99 days °C^−1^ (temporal). Field observation estimated sensitivities of −2.96 days °C^−1^ with a standard deviation of 2.74 days °C^−1^ (spatial) and −4.77 days °C^−1^ with a standard deviation of 3.22 days °C^−1^ (temporal). Out of 74 species in the herbarium data and 32 species in the field observations, 24 species were shared between these two datasets. For spatial sensitivity, 16 out of 24 (67%) species showed consistency between the two datasets. There was a tendency of decreasing spatial sensitivity: 16 species (67%) showed a short-term sensitivity from field observation weaker than long-term sensitivity from herbarium, with 7 species (29%) significant. For temporal sensitivity, 20 out of 24 species (83%) showed consistency between the two datasets. There was a tendency for increasing temporal sensitivity: 17 species (71%) showed a short-term sensitivity from field observation stronger than long-term sensitivity from herbarium, but with 3 species (13%) significant.

**Figure 3.**
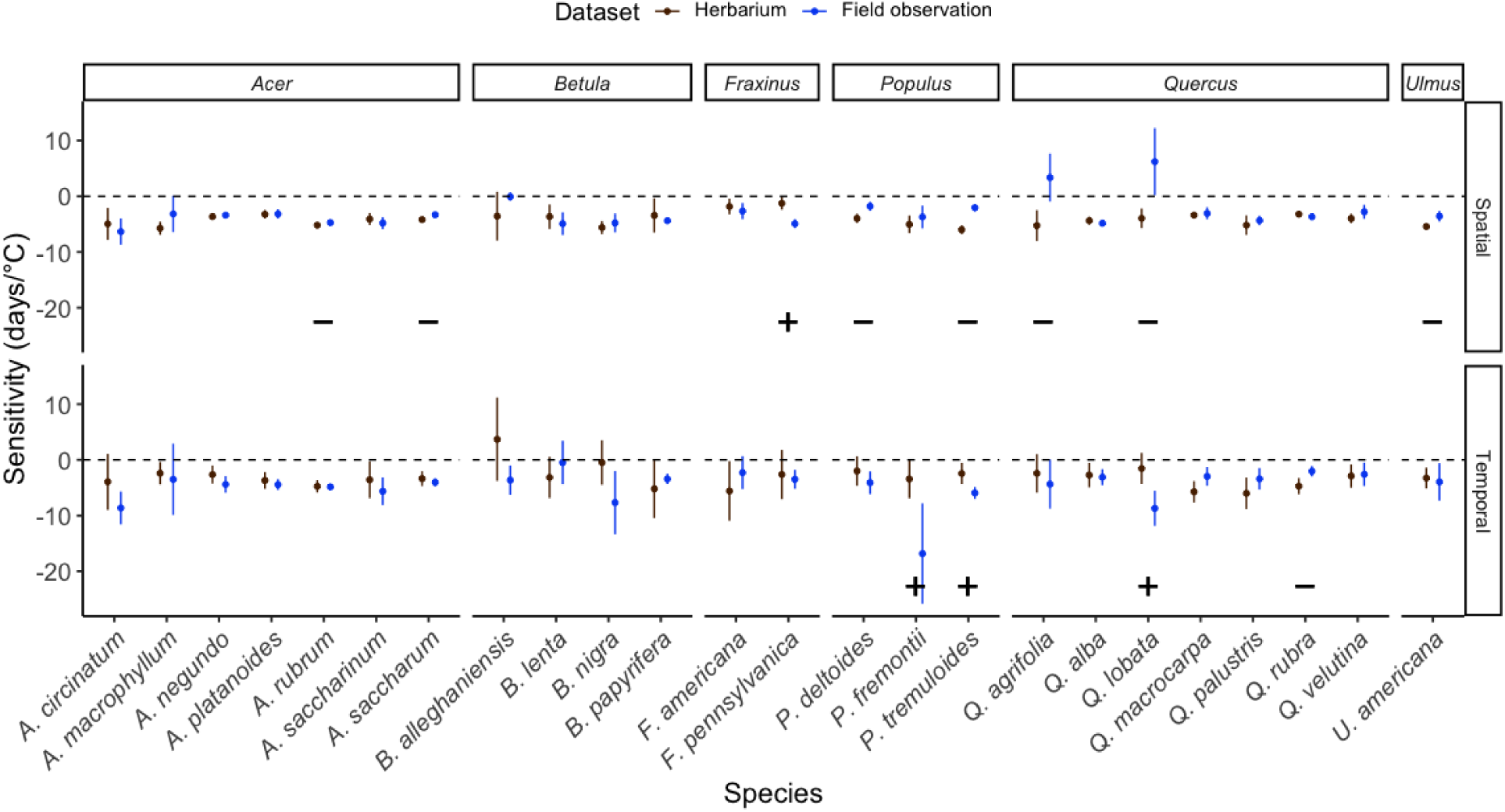
Comparison of spatial and temporal sensitivity estimated from the long- and short-term datasets. Symbol “+” represents a significantly stronger sensitivity over short-term estimates compared with long-term estimates, represented by a significantly stronger (more negative) sensitivity derived from short-term field observations compared to that from long-term herbarium data. Symbol “−” represents a significantly weaker sensitivity over short-term estimates compared with long-term estimates, both at 99% confidence intervals.

### 3.3 Strong phylogenetic signals in spatial, not temporal, sensitivity

The standard mixed-effects model and phylogenetic mixed-effects models yielded similar estimates for both the mean and standard deviation of spatial and temporal sensitivity. The overall spatial sensitivity across species was −4.06 days °C^−1^ (from SMM) or −4.00 days °C^−1^(from PMM), indicating 4 days advancement in wind-pollinated trees’ flowering time for every 1°C temperature increase over space. Species differed in spatial sensitivities with a 95% credible interval of −4.47 to −3.65 days °C^−1^ (from SMM) or −5.69 to −2.40 days °C^−1^ (from PMM). For temporal sensitivity, the overall estimate across species was −2.87 days °C^−1^ (from SMM) or −2.84 days °C^−1^ (from PMM), indicating that 3 days of advancement in wind-pollinated trees’ flowering time for every 1°C temperature increase over time. Species differed in temporal sensitivities with a 95% credible interval of −3.68 to −2.04 days °C^−1^ (from SMM) or −4.12 to −1.36 days °C^−1^ (from PMM).

Spatial sensitivity exhibited a strong phylogenetic signal (Pagel’s *λ* > 0.5; Pearse et al., 2025), with a Pagel’s *λ* value of 0.74 (95% credible interval: 0.21–0.99). The strong phylogenetic signal of spatial sensitivity was evident in the phylogenetic tree, where closely related species tended to display similar levels of spatial sensitivity (Figure 4). Among the genera, *Fraxinus* tended to have weaker-than-average spatial sensitivities, while *Ulmus* exhibited stronger-than-average spatial sensitivities. In contrast, temporal sensitivity demonstrated only a weak phylogenetic signal (Pagel’s *λ* = 0.35; 95% credible interval: 0.01–0.91, Figure S2).

**Figure 4.**
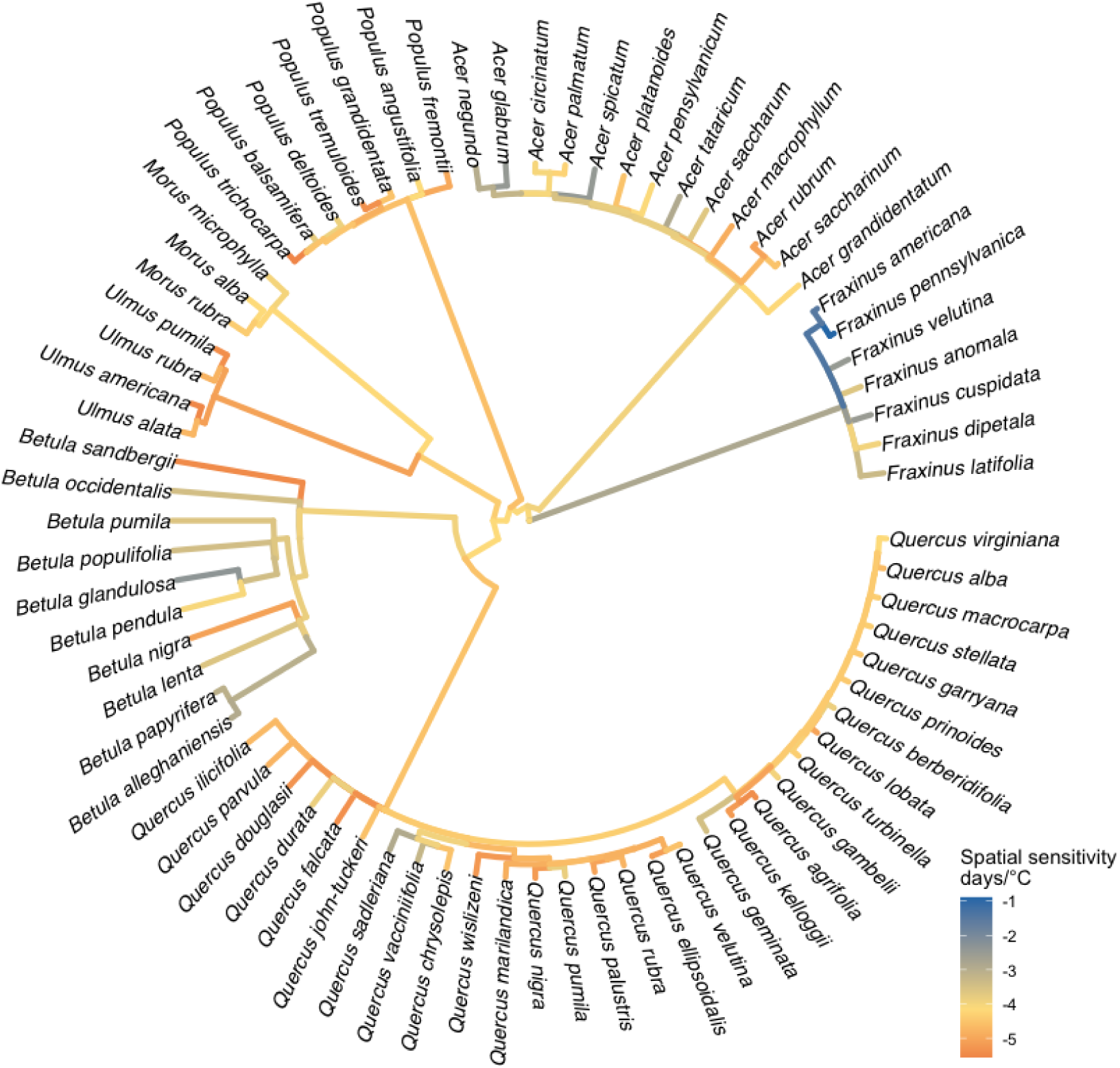
Species-level spatial sensitivity from phylogenetic mixed-effects models. The phylogenetic tree with color indicating the sensitivity of the corresponding leaves in the phylogenetic mixed-effects model. Yellow is the mean sensitivity across species. Blue indicates weaker sensitivity than the mean, while red indicates stronger.

## 4. Discussion

To understand the flowering phenology responses of wind-pollinated trees to temperature changes, we estimated spatial and temporal sensitivities across 74 species on the continental-US scale. Spatial sensitivities are around −4.00 to −4.89 days °C^−1^, and temporal sensitivities are about −2.84 to −3.68 days °C^−1^. We evaluated the role of adaptation, time scale, and interspecific variations in the sensitivities. We found that there is no significant difference between spatial and temporal sensitivity for 77% of the species, indicating that plasticity dominated wind-pollinated trees’ flowering time sensitivity to temperature. Most species showed consistent responses between the last century and the most recent two decades. The variations of sensitivity among species showed strong phylogenetic signals for spatial sensitivity.

### 4.1 Dominance of plasticity in the temperature sensitivity of flowering time in wind-pollinated trees

We found plasticity dominated the temperature sensitivity of flowering time for 77% of the wind-pollinated tree species we analyzed, despite an overall tendency for co-gradient adaptation (Table 1). Though this weak impact of adaptation compared to plasticity has been reported in common garden experiments (Firmat et al., 2017; Guo et al., 2023; McDonough MacKenzie et al., 2018; Zeng & Wolkovich, 2024), our analysis provides a quantitative evaluation of the magnitude of plasticity and adaptation in the form of sensitivity with a broad spatial, temporal, and taxonomy coverage. The general consistency between spatial and temporal sensitivities generally supports the use of space-for-time substitution to estimate future flowering times for most wind-pollinated trees (Buyantuyev et al., 2012), with a potential bias toward overestimating temporal changes. We also identified a few species that might need caution when applying space-for-time substitution, such as *Quercus nigra*.

The limited adaptation observed in wind-pollinated trees can be attributed to their life-history traits. Our findings, although qualitatively consistent with previous findings on grass and shrub species (Ramirez-Parada et al., 2024), suggested plasticity might play an even more important role in wind-pollinated trees, with 77% of the species showing no significant differences between spatial and temporal sensitivity (41% in Ramirez-Parada et al., 2024). This result likely reflects fundamental life-history constraints: species with shorter life spans can undergo evolutionary changes more rapidly due to their shorter generation times, whereas long-lived species such as trees might rely more on plasticity to respond to environmental variability (Cordero & Janzen, 2013). The limited evidence for significant local adaptation in long-lived wind-pollinated trees indicates that these trees may face greater vulnerability to climate change. Their slower rate of molecular evolution might limit their ability to keep pace with rapidly changing environmental conditions (Smith & Beaulieu, 2009).

Though dominated by plasticity, we observed a tendency of co-gradient adaptation, with 50 out of 74 (68%) species showing a spatial sensitivity stronger than temporal sensitivity (12 significant and 38 non-significant, Figure 2). A similar tendency of co-gradient adaptation was reported for grasses and shrubs (Ramirez-Parada et al., 2024). The observed co-gradient adaptation suggests that local adaptation tends to accentuate, rather than diminish, flowering time differences along latitudinal gradients (Love & Mazer, 2021). One possible explanation is that phenotypic plasticity is constrained by legacy effects, wherein environmental conditions from the preceding year influence a plant’s phenological responses in the subsequent growing season. These legacy effects, which are particularly evident in temporal sensitivity, may limit the ability of plants to fully respond to current temperature conditions (Mulder et al., 2017). However, local adaptation to consistently warm or cold environments might enhance the plant’s capacity to adjust to prevailing climatic conditions.

### 4.2 No significant difference between short- and long-term sensitivity

Most species showed consistent sensitivity estimates when comparing the century-long herbarium records with the two-decade field observations. Consistency was observed for 16 out of 24 species (67%) in spatial sensitivities and for 20 out of 24 species (83%) in temporal sensitivities (Figure 3, Tables S1 & S2). These results suggest that the previously reported decline in temperature sensitivity (Fu et al., 2015; Güsewell et al., 2017; Xiong et al., 2024) is not evident in the wind-pollinated trees we studied, prompting us to reconsider the mechanisms commonly proposed to explain this decrease in sensitivity. The first proposed mechanism attributes declining sensitivity to limitations imposed by other environmental cues, such as photoperiod and water availability, which may constrain the response of flowering time to temperature increases (Fu et al., 2015; Xiong et al., 2024). Our results suggested that the flowering phenology of the wind-pollinated trees we studied has not yet been limited by non-thermal cues, which might explain stronger temperature sensitivities of wind-pollinated trees compared to herbaceous species and other pollination types (Calinger et al., 2013). A second prevailing explanation for the decline in sensitivity centers on the nonlinear nature of phenological responses to temperature, whereby continued warming yields diminishing marginal effects on flowering advancement, leading to a decline in observed linear sensitivity over time (Wolkovich et al., 2021). While theoretically highly plausible, our results suggested that nonlinear phenological changes are not yet evident among our species of interest within the range of temperatures they experienced in the field. Strongly linear responses have also been shown in a whole-ecosystem warming experiment, with 0°C to 9°C warming (Richardson et al., 2018). The third current explanation involves adaptation, as previously discussed: long-lived trees may require much more time to exhibit detectable adaptive responses to climate change.

### 4.3 Flowering time temperature sensitivity across species variation and the phylogenetic signal

The PMM model reports a spatial sensitivity of −4.00 (95% credible interval: −5.69 to −2.40) days °C^−1^ and a temporal sensitivity of −2.84 (95% credible interval: −4.12 to −1.36) days °C^−1^. These estimates align with previously reported ranges of temperature sensitivity (Geng et al., 2022; Wang et al., 2020), but we further differentiate between spatial and temporal temperature sensitivities. Spatial sensitivity exhibits a significant phylogenetic signal (Figure 4), indicating that interspecific differences in spatial sensitivity are associated with evolutionary relationships; closely related species tend to exhibit similar responses. *Fraxinus* generally showed lower-than-average spatial sensitivities, whereas *Ulmus* demonstrated higher-than-average spatial sensitivities. This variation in spatial sensitivity may reflect differences in the relative timing or even the sequence of flowering between these genera along environmental gradients. That is, the flowering time gap between *Fraxinus* and *Ulmus* may vary by location. In contrast, temporal sensitivity showed a weak phylogenetic signal, with species exhibiting similar sensitivities. This implies that, within the same location, different species adjust their flowering time in response to temperature changes at comparable rates, which contrasts with predictions of shifts in the order of spring phenology in temperate forests (Roberts et al., 2015). The presence of a phylogenetic signal may occur because spatial sensitivity reflects both phenotypic plasticity and local adaptation, with the latter shaped by evolutionary history (Morales-Castilla et al., 2024).

### 4.4 Limitations and future directions

While our analysis effectively estimated spatial and temporal sensitivity across broad geographic and temporal scales, a key limitation lies in the greater uncertainty associated with temporal sensitivity estimates, likely driven by the narrower temperature variation available for inference. Our analysis shows high uncertainty in the temporal sensitivity estimates, which is reflected in the wider confidence intervals compared to those for spatial sensitivity estimates (Figure 2). This uncertainty in temporal sensitivity could have partly contributed to challenges in detecting significant differences in spatial sensitivities. This uncertainty could be attributed to the narrow range of temperature variation used to infer temporal sensitivity, which was limited to −2°C to 2°C, leading to a low signal-to-noise ratio. In contrast, the inference of spatial sensitivity typically benefits from a much broader temperature range, from 0°C to 25°C (Figure 1). Future studies are likely to provide more precise and reliable estimates of temporal sensitivity through longer time series, controlled experiments, and attention to years with temperature anomalies.

Our findings provide actionable insights for phenological forecasting under climate change by identifying dominant plastic responses, stable sensitivities across time, and strong phylogenetic structuring as key patterns shaping flowering behavior in wind-pollinated trees. The observed consistency between spatial and temporal sensitivities validates the use of the space-for-time substitution method for projecting future flowering times (Buyantuyev et al., 2012). We also identified a subset of species for which this approach warrants caution, as their spatial and temporal sensitivities differ markedly, likely reflecting pronounced local adaptation. The demonstrated stability of sensitivity over time supports the reliability of using short-term sensitivity measurements to make long-term, at least over the scale of a century, phenological forecasts for trees that respond linearly to temperature change (Richardson et al., 2018). The strong phylogenetic signal detected in spatial sensitivity suggests that incorporating phylogenetic relationships can improve predictions of climate-driven flowering responses, particularly for poorly sampled species within well-sampled clades (Morales-Castilla et al., 2024).

## 5. Conclusions

This study assessed how the flowering time in 74 wind-pollinated tree species responds to temperature changes by systematically analyzing spatial and temporal trends over both the past century and the last two decades. The sensitivity is generally around −2.84 to −4.89 days °C^−1^, indicating that every Celsius temperature increase leads to 2.84 to 4.89 days advance in flowering time. We found limited evidence for local adaptation, with 77% of species showing plasticity-dominated responses, possibly due to the long lifespan of trees. Our comparison of long-term herbarium and short-term field data showed consistent sensitivity estimates over the last century and the last two decades for wind-pollinated trees, in contrast to the previously observed decrease in temperature sensitivity in recent years. Lastly, we detected a strong phylogenetic signal in spatial sensitivity, suggesting that closely related species exhibit similar spatial sensitivities. Overall, our findings underscore the dominant role of plasticity, the stability of sensitivity over time, and the importance of phylogenetic structure for the flowering phenology of wind-pollinated trees. These insights can improve phenological forecasting under climate change.

## Author contributions

Yi Liu: Conceptualization, Methodology, Data Curation, Formal Analysis, Visualization, Writing – Original Draft, Writing – Review & Editing, Project Administration

Yiluan Song: Conceptualization, Methodology, Validation, Writing – Review & Editing

William Weaver: Methodology, Data Curation, Validation

Kerby Shedden: Methodology, Validation

Yang Chen: Methodology, Validation

Stephen Smith: Methodology, Writing – Review & Editing, Validation

Kai Zhu: Conceptualization, Resources, Writing – Review & Editing, Supervision

## Data availability statement

All primary data are publicly available, and all generated data will be permanently archived upon acceptance. Code is available on GitHub: https://github.com/zhulabgroup/phenology-sensitivity/tree/clean and will be archived upon acceptance.

## Acknowledgments

We thank Ignacio Morales-Castilla for providing code from their paper (Morales-Castilla et al., 2024), which was helpful in building both the standard mixed-effects model and the phylogenetic mixed-effects model. We thank members of Dr. Kai Zhu’s and Dr. Yang Chen’s research groups at the University of Michigan for feedback. This research is supported by the National Science Foundation (NSF) grant 2306198 and the Michigan Institute for Data and AI in Society (MIDAS) Propelling Original Data Science (PODS) grant in 2023.

## References

Anderegg, W. R. L., Abatzoglou, J. T., Anderegg, L. D. L., Bielory, L., Kinney, P. L., & Ziska, L. (2021). Anthropogenic climate change is worsening North American pollen seasons. Proceedings of the National Academy of Sciences, 118(7), e2013284118. 10.1073/pnas.2013284118

Blois, J. L., Williams, J. W., Fitzpatrick, M. C., Jackson, S. T., & Ferrier, S. (2013). Space can substitute for time in predicting climate-change effects on biodiversity. Proceedings of the National Academy of Sciences, 110(23), 9374–9379. 10.1073/pnas.1220228110

Blumstein, M., Webster, S., Hopkins, R., Basler, D., Yun, J., & Des Marais, D. L. (2025). Genomics highlight an underestimation of phenology sensitivity to the urban heat island effect. Proceedings of the National Academy of Sciences, 122(12), e2408564122. 10.1073/pnas.2408564122

Calinger, K. M., Queenborough, S., & Curtis, P. S. (2013). Herbarium specimens reveal the footprint of climate change on flowering trends across north-central North America. Ecology Letters, 16(8), 1037–1044. 10.1111/ele.12135

Cordero, G., & Janzen, F. (2013). Does life history affect molecular evolutionary rates?

Davies, T. J., Wolkovich, E. M., Kraft, N. J. B., Salamin, N., Allen, J. M., Ault, T. R., Betancourt, J. L., Bolmgren, K., Cleland, E. E., Cook, B. I., Crimmins, T. M., Mazer, S. J., McCabe, G. J., Pau, S., Regetz, J., Schwartz, M. D., & Travers, S. E. (2013). Phylogenetic conservatism in plant phenology. Journal of Ecology, 101(6), 1520–1530. 10.1111/1365-2745.12154

Evans, M. E. K., Adler, P. B., Angert, A. L., Dey, S. M. N., Girardin, M. P., Heilman, K. A., Klesse, S., Perret, D. L., Sax, D. F., Sheth, S. N., Stemkovski, M., & Williams, J. L. (2025). Reconsidering space-for-time substitution in climate change ecology. Nature Climate Change, 15(8), 809–812. 10.1038/s41558-025-02392-0

Firmat, C., Delzon, S., Louvet, J.-M., Parmentier, J., & Kremer, A. (2017). Evolutionary dynamics of the leaf phenological cycle in an oak metapopulation along an elevation gradient. Journal of Evolutionary Biology, 30(12), 2116–2131. 10.1111/jeb.13185

Fitter, A. H., & Fitter, R. S. R. (2002). Rapid Changes in Flowering Time in British Plants. Science, 296(5573), 1689–1691. 10.1126/science.1071617

Fox, R. J., Donelson, J. M., Schunter, C., Ravasi, T., & Gaitán-Espitia, J. D. (2019). Beyond buying time: The role of plasticity in phenotypic adaptation to rapid environmental change. Philosophical Transactions of the Royal Society B: Biological Sciences, 374(1768), 20180174. 10.1098/rstb.2018.0174

Fu, Y. H., Zhao, H., Piao, S., Peaucelle, M., Peng, S., Zhou, G., Ciais, P., Huang, M., Menzel, A., Peñuelas, J., Song, Y., Vitasse, Y., Zeng, Z., & Janssens, I. A. (2015). Declining global warming effects on the phenology of spring leaf unfolding. Nature, 526(7571), 104–107. 10.1038/nature15402

GBIF.org (13 May 2024) GBIF Occurrence Download 10.15468/dl.uk96qu

Geng, X., Fu, Y. H., Piao, S., Hao, F., De Boeck, H. J., Zhang, X., Chen, S., Guo, Y., Prevéy, J. S., Vitasse, Y., Peñuelas, J., Janssens, I. A., & Stenseth, N. Chr. (2022). Higher temperature sensitivity of flowering than leaf-out alters the time between phenophases across temperate tree species. Global Ecology and Biogeography, 31(5), 901–911. 10.1111/geb.13463

Guo, X., Buttò, V., Mohytych, V., Klisz, M., Surget-Groba, Y., Huang, J., Delagrange, S., & Rossi, S. (2023). Plasticity plays a dominant role in regulating the phenological variations of sugar maple populations in Canada. Frontiers in Ecology and Evolution, 11. 10.3389/fevo.2023.1217871

Güsewell, S., Furrer, R., Gehrig, R., & Pietragalla, B. (2017). Changes in temperature sensitivity of spring phenology with recent climate warming in Switzerland are related to shifts of the preseason. Global Change Biology, 23(12), 5189–5202. 10.1111/gcb.13781

Knapp, A. K., Condon, K. V., Folks, C. C., Sturchio, M. A., Griffin-Nolan, R. J., Kannenberg, S. A., Gill, A. S., Hajek, O. L., Siggers, J. A., & Smith, M. D. (2024). Field experiments have enhanced our understanding of drought impacts on terrestrial ecosystems—But where do we go from here? Functional Ecology, 38(1), 76–97. 10.1111/1365-2435.14460

Knott, J. A., Liang, L., Dukes, J. S., Swihart, R. K., & Fei, S. (2023). Phenological response to climate variation in a northern red oak plantation: Links to survival and productivity. Ecology, 104(3), e3940. 10.1002/ecy.3940

Kudo, G., & Ida, T. Y. (2013). Early onset of spring increases the phenological mismatch between plants and pollinators. Ecology, 94(10), 2311–2320. 10.1890/12-2003.1

Lo, F., Bitz, C. M., Battisti, D. S., & Hess, J. J. (2019). Pollen calendars and maps of allergenic pollen in North America. Aerobiologia, 35(4), 613–633. 10.1007/s10453-019-09601-2

Love, N. L. R., & Mazer, S. J. (2021). Region-specific phenological sensitivities and rates of climate warming generate divergent temporal shifts in flowering date across a species’ range. American Journal of Botany, 108(10), 1873–1888. 10.1002/ajb2.1748

McDonough MacKenzie, C., Primack, R. B., & Miller-Rushing, A. J. (2018). Local environment, not local adaptation, drives leaf-out phenology in common gardens along an elevational gradient in Acadia National Park, Maine. American Journal of Botany, 105(6), 986–995. 10.1002/ajb2.1108

Menzel, A., Sparks, T. H., Estrella, N., Koch, E., Aasa, A., Ahas, R., Alm-Kübler, K., Bissolli, P., Braslavská, O., Briede, A., Chmielewski, F. M., Crepinsek, Z., Curnel, Y., Dahl, Å., Defila, C., Donnelly, A., Filella, Y., Jatczak, K., Måge, F., … Zust, A. (2006). European phenological response to climate change matches the warming pattern. Global Change Biology, 12(10), 1969–1976. 10.1111/j.1365-2486.2006.01193.x

Morales-Castilla, I., Davies, T. J., Legault, G., Buonaiuto, D. M., Chamberlain, C. J., Ettinger, A. K., Garner, M., Jones, F. A. M., Loughnan, D., Pearse, W. D., Sodhi, D. S., & Wolkovich, E. M. (2024). Phylogenetic estimates of species-level phenology improve ecological forecasting. Nature Climate Change, 14(9), 989–995. 10.1038/s41558-024-02102-2

Mulder, C. P. H., Iles, D. T., & Rockwell, R. F. (2017). Increased variance in temperature and lag effects alter phenological responses to rapid warming in a subarctic plant community. Global Change Biology, 23(2), 801–814. 10.1111/gcb.13386

Ng, A. E., & Boersma, P. (2023). Diagnosed Allergic Conditions in Adults: United States, 2021. NCHS Data Brief, 460, 1–8.

Perret, D. L., Evans, M. E. K., & Sax, D. F. (2024). A species’ response to spatial climatic variation does not predict its response to climate change. Proceedings of the National Academy of Sciences, 121(1), e2304404120. 10.1073/pnas.2304404120

Primack, R. B., Vaughn, S., & Terry, C. (2025). Local soil temperature advances flowering phenology of Canada mayflower (Maianthemum canadense), with implications for climate change assessment. Oecologia, 207(2), 36. 10.1007/s00442-025-05668-6

Ramirez-Parada, T. H., Park, I. W., Record, S., Davis, C. C., Ellison, A. M., & Mazer, S. J. (2024). Plasticity and not adaptation is the primary source of temperature-mediated variation in flowering phenology in North America. Nature Ecology & Evolution, 1–10. 10.1038/s41559-023-02304-5

Richardson, A. D., Hufkens, K., Milliman, T., Aubrecht, D. M., Furze, M. E., Seyednasrollah, B., Krassovski, M. B., Latimer, J. M., Nettles, W. R., Heiderman, R. R., Warren, J. M., & Hanson, P. J. (2018). Ecosystem warming extends vegetation activity but heightens vulnerability to cold temperatures. Nature, 560(7718), 368–371. 10.1038/s41586-018-0399-1

Roberts, A. M. I., Tansey, C., Smithers, R. J., & Phillimore, A. B. (2015). Predicting a change in the order of spring phenology in temperate forests. Global Change Biology, 21(7), 2603–2611. 10.1111/gcb.12896

Rosemartin, A. H., Denny, E. G., Gerst, K. L., Marsh, R. L., Posthumus, E. E., Crimmins, T. M., & Weltzin, J. (2018). USA National Phenology Network observational data documentation. In Open-File Report (Nos. 2018–1060). U.S. Geological Survey. 10.3133/ofr20181060

Smith, S. A., & Beaulieu, J. M. (2009). Life history influences rates of climatic niche evolution in flowering plants. Proceedings of the Royal Society B: Biological Sciences, 276(1677), 4345–4352. 10.1098/rspb.2009.1176

Valdés, A., Arnold, P. A., & Ehrlén, J. (2023). Spring temperature drives phenotypic selection on plasticity of flowering time. Proceedings of the Royal Society B: Biological Sciences, 290(2006), 20230670. 10.1098/rspb.2023.0670

Wang, H., Wang, H., Ge, Q., & Dai, J. (2020). The Interactive Effects of Chilling, Photoperiod, and Forcing Temperature on Flowering Phenology of Temperate Woody Plants. Frontiers in Plant Science, 11, 443. 10.3389/fpls.2020.00443

Weaver, W. N., & Smith, S. A. (2023). From leaves to labels: Building modular machine learning networks for rapid herbarium specimen analysis with LeafMachine2. Applications in Plant Sciences, 11(5), e11548. 10.1002/aps3.11548

Wolkovich, E. M., Auerbach, J., Chamberlain, C. J., Buonaiuto, D. M., Ettinger, A. K., Morales-Castilla, I., & Gelman, A. (2021). A simple explanation for declining temperature sensitivity with warming. Global Change Biology, 27(20), 4947–4949. 10.1111/gcb.15746

Wolkovich, E. M., Cook, B. I., Allen, J. M., Crimmins, T. M., Betancourt, J. L., Travers, S. E., Pau, S., Regetz, J., Davies, T. J., Kraft, N. J. B., Ault, T. R., Bolmgren, K., Mazer, S. J., McCabe, G. J., McGill, B. J., Parmesan, C., Salamin, N., Schwartz, M. D., & Cleland, E. E. (2012). Warming experiments underpredict plant phenological responses to climate change. Nature, 485(7399), Article 7399. 10.1038/nature11014

Xie, Y., Thammavong, H. T., & Park, D. S. (2022). The ecological implications of intra- and inter-species variation in phenological sensitivity. New Phytologist, 236(2), 760–773. 10.1111/nph.18361

Xiong, T., Du, S., Zhang, H., & Zhang, X. (2024). Decreasing temperature sensitivity of spring phenology decelerates the advance of spring phenology in northern temperate and boreal forests. Ecological Indicators, 161, 111983. 10.1016/j.ecolind.2024.111983

Zablotsky, B., Black, L. I., & Akinbami, L. J. (2023). Diagnosed Allergic Conditions in Children Aged 0-17 Years: United States, 2021. NCHS Data Brief, 459, 1–8.

Zeng, Z. A., & Wolkovich, E. M. (2024). Weak evidence of provenance effects in spring phenology across Europe and North America. New Phytologist, 242(5), 1957–1964. 10.1111/nph.19674

Ziello, C., Böck, A., Estrella, N., Ankerst, D., & Menzel, A. (2012). First flowering of wind-pollinated species with the greatest phenological advances in Europe. Ecography, 35(11), 1017–1023. 10.1111/j.1600-0587.2012.07607.x

