## Supplementary materials for "Dominance of plasticity in wind-pollinated trees’ flowering time response to temperature"

### Supplementary Methods:

#### Supplementary Method 1:

Robust linear regression. Since the dataset contains outliers, we used robust linear regression to mitigate their influence by iteratively fitting and adjusting observation weights in the optimization process. We used the *rlm* function in the *MASS* package with M-estimation and Huber weighting. The M-estimation defines the equation to be optimized as the weighted square residuals and uses the first derivative equals to 0 to calculate the optimized parameters (Equation 1). Huber weighting defines the reweight rule: the weight is 1 if the residual is smaller than the threshold *k*, and the weight decreases with the residual when the residual is larger than *k* (Equation 2):

$$\mathbf{b}_j = (\mathbf{X}'\mathbf{W}_{j-1}\mathbf{X})^{-1}\mathbf{X}'\mathbf{W}_{j-1}\mathbf{Y} \quad (1)$$

$$w(r) = \begin{cases} 1 & \text{if } |r| \le k \\ \frac{k}{|r|} & \text{if } |r| > k \end{cases} \quad (2)$$

where $j$ is the iteration indicator, $\boldsymbol{b}$ is the coefficient vector of the linear models, $w(r)$ is the weight of observation with residual $r$. $k$ is the threshold for weight adjustment. We used the default value $1.345\sigma$, where $\sigma$ is the residual standard deviation. Model fitting starts with all weights set to 1. After the initial fit, observations will be reweighted based on residuals to refit the model. This process is repeated until changes become negligible (convergence is reached). We used 30 interactions to make sure the model converged for each species. Also, to ensure robustness estimates of the parameters, we require at least 10 distinct values for $d$, $T_{\mathrm{norm}}$, and $T_{\mathrm{anom}}$, and a minimum of 30 unique combinations for a species to be included. We also verified the correlation between $T_{\mathrm{norm}}$ and $T_{\mathrm{anom}}$ for each species to be below 0.5.

#### Supplementary Method 2:

Wald test. We employed the Wald test to assess whether the two coefficients ($\beta_{\text{spatial}}$ and $\beta_{\text{temporal}}$) in the fitted model differ significantly from each other. Under the null hypothesis that $\beta_{\text{spatial}} = \beta_{\text{temporal}}$, the test statistic (computed in Equations 3 and 4) follows a $\chi^2$-distribution with 1 degree of freedom. A p-value < 0.05 provides evidence to reject the null hypothesis.

$$\mathrm{Var}(\hat{\beta}_{\text{spatial}} - \hat{\beta}_{\text{temporal}}) = \mathrm{Var}(\hat{\beta}_{\text{spatial}}) + \mathrm{Var}(\hat{\beta}_{\text{temporal}}) - 2 \cdot \mathrm{Cov}(\hat{\beta}_{\text{spatial}}, \hat{\beta}_{\text{temporal}}) \quad (3)$$

$$\chi^2 = \frac{(\hat{\beta}_{\text{spatial}} - \hat{\beta}_{\text{temporal}})^2}{\mathrm{Var}(\hat{\beta}_{\text{spatial}} - \hat{\beta}_{\text{temporal}})} \quad (4)$$

#### Supplementary Method 3: Fitting hierarchical models in Stan

Standard mixed-effects model (SMM). We used the normal distribution as the prior for $\mu$ and the inverse gamma distribution for $\sigma$. Posterior distributions were obtained using the No-U-Turn sampler variant of Hamiltonian Monte Carlo in RStan as implemented in R v.4.4.2 using the RStan package v.2.32.7. Sampling was done using four Markov chain Monte Carlo (MCMC) chains with 4,000 iterations and a warmup of 2,000 for each. All estimates had Gelman–Rubin statistics (R-hat) between 0.9995 and 1.0021 and a minimum effective sample size of 1,491. Over 75% of parameters have an effective sample size exceeding 14,000.

Phylogenetic mixed-effects model (PMM). We used the normal distribution as the prior for $\mu$, the inverse gamma distribution for $\sigma$, and the beta distribution for $\lambda$. Posterior distributions were obtained using the same methods and software as HMM. Sampling was done using four MCMC chains with 8,000 iterations and a warmup of 4,000 for each. All estimates had Gelman–Rubin statistics (R-hat) between 0.9998 and 1.0027 and a minimum effective sample size of 1,748. Over 75% of parameters have an effective sample size exceeding 19,000.

### Supplementary Figures:

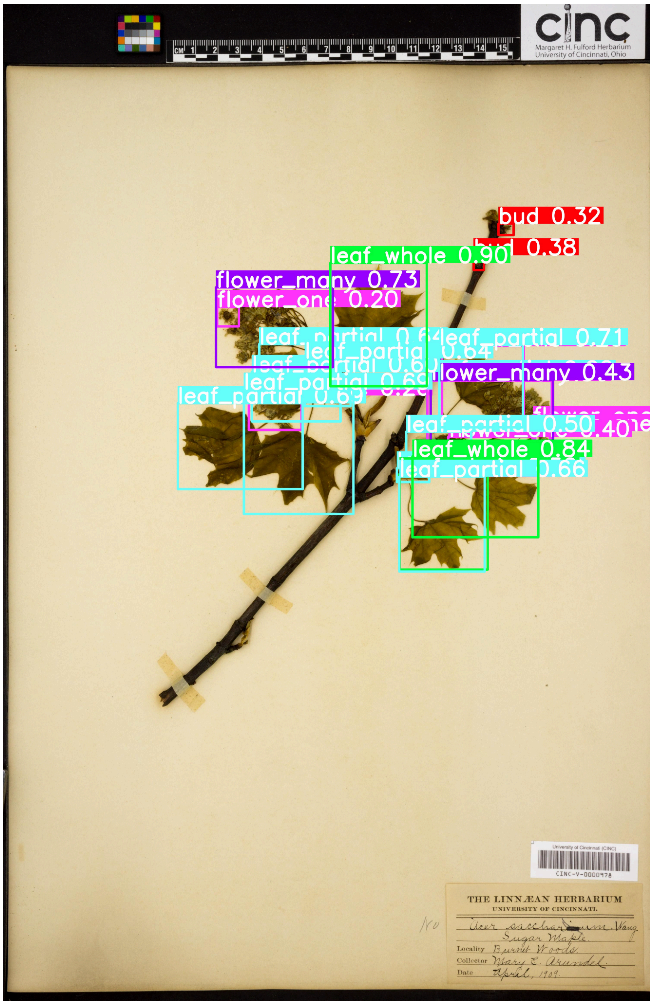


**Supplementary Figure 1.** An image of an *Acer saccharum* (sugar maple) specimen collected in 1909 (downloaded from GBIF). The colored squares are the output of LeafMachine2, showing the detection and classification of the part. The number shows the confidence level (between zero and one), with a higher value indicating higher confidence.

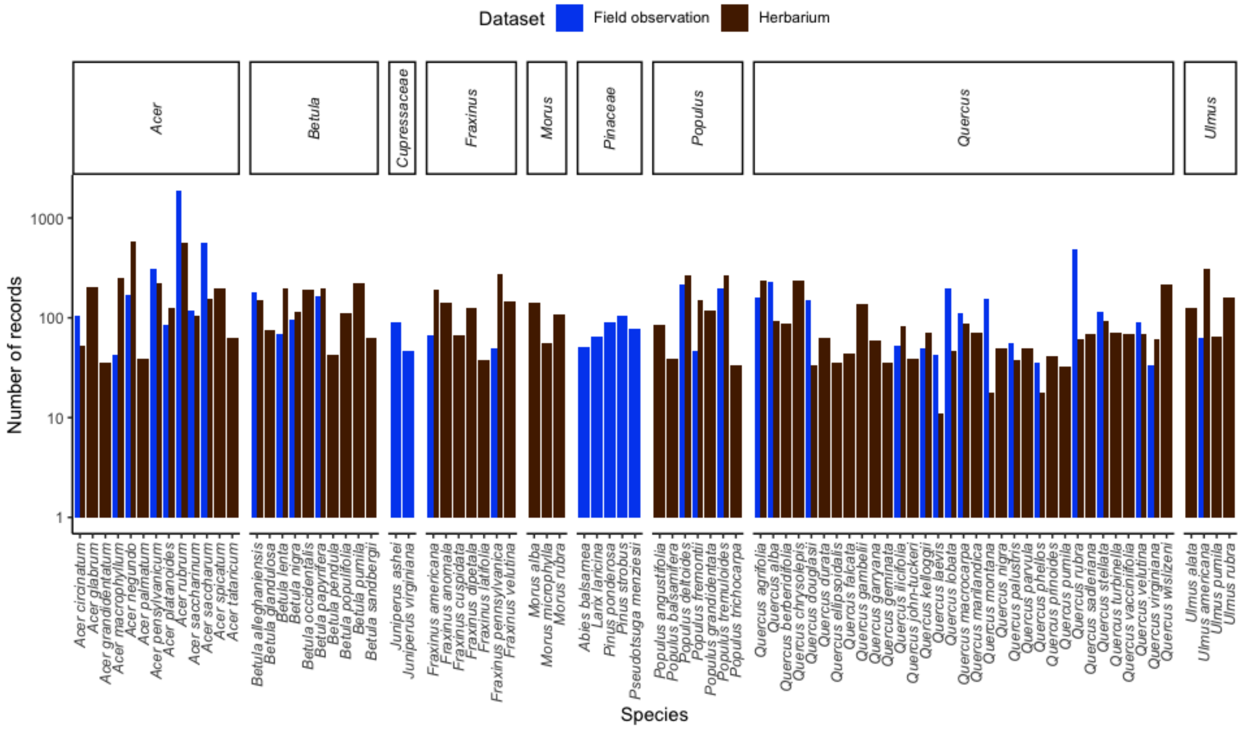


**Supplementary Figure 2.** Sample size of each species in field observations and herbarium. The y-axis is in log scale.
